# HP2NET: Empowering Efficient Phylogenetic Network Analysis through High-Performance Computing

**DOI:** 10.64898/2026.03.05.709005

**Authors:** Rafael Terra, Diego Carvalho, Denis Jacob Machado, Carla Osthoff, Kary Ocaña

## Abstract

Advances in High-Performance Computing (HPC) have enabled increasingly complex genomic analyses, including those in phylogenomics. These analyses contribute to understanding the evolution of viruses and pathogens, improving our knowledge of disease transmission, and supporting targeted public health strategies. However, due to the increasing number of tools and processing steps involved, executing these analyses manually, step by step, becomes error-prone and inefficient. To address this challenge, we present HP2NET, a robust framework for reproducible, efficient, and scalable phylogenetic network analysis. HP2NET integrates five workflows based on state-of-the-art tools such as PhyloNetworks and PhyloNet, allowing the analysis of multiple datasets and workflows in a single execution. The framework includes features such as task packaging and data reuse to improve performance and resource utilization in HPC environments. We perform a comprehensive performance evaluation of the software used within HP2NET, identifying bottlenecks and analyzing gains from parallel processing. Data reuse provided up to 15.35% time reduction, for a small dataset, in our experimental environment, while parallel execution of the five pipelines reduced total runtime by up to 90.96% compared to sequential runs. Finally, we validate HP2NET in a real-world case study by analyzing Dengue virus genomes, demonstrating its applicability value for large-scale phylogenetic analyses.

## I. Introduction

THE landscape of genomics research drives significant advancements in high-throughput DNA sequencing, supported by high-performance computing (HPC). This transformation enhances our understanding of life sciences and means to fight diseases [1], [2]. For example, the National Institutes of Health (NIH) HPC system integrates robust data management capabilities into its scientific applications, enabling researchers to process and analyze large-scale genomic data to extract relevant information. The NIH-Wide Strategic Plan for COVID-19 Research has significantly contributed to the global pandemic response [3]. Additionally, the role of viral genomics in fighting outbreaks of Zika virus (ZIKV) in Latin America,

Yellow fever virus (YFV) in Angola and Brazil, West Nile virus (WNV) in the Americas, Chikungunya virus (CHIKV) in East Africa, and the ongoing Dengue virus (DENV) pandemic in the tropics and subtropics underscore the critical role of HPC to make sense of viral genomes [4].

HPC-accelerated phylogenetic methods provide critical insights into the evolution of viruses and pathogens, improving our understanding of disease spread and control and enhancing health outcomes through targeted public health strategies [5]. Building on these advancements, phylogenomics workflows [1], [2] can be integrated into scientific workflow management systems (SWfMSs) such as Pegasus [6] and NextFlow [7]. These systems, alongside language-parallel scripting libraries such as Swift [8] or Parsl [9], enhance workflow management, improving efficiency and scalability in phylogenomics, especially in cloud environments and HPC clusters [10], [11].

Despite the growing capabilities of HPC and SWfMSs, the vast network space makes the reconstruction of phylogenetic networks challenging in phylogenetics and genome evolution. One approach is solving the minimum phylogenetic network problem, where phylogenetic trees are first inferred, and then the smallest network displaying all the trees is computed [12]. The most significant limitation of methods to infer introgression and hybridization, including species network methods, is scalability. These methods have only been used with a handful of taxa and fewer than 200 loci [13]. Recent innovations incorporate reticulate evolutionary processes and reconstruct networks from ancestral profiles to improve phylogenetic network analyses [14], [15].

Effective modeling of scientific workflows, particularly in large-scale phylogenomics analyses, requires integrating several software programs, and dataflows and provenance management are particularly important. For example, [16] introduces a workflow for phylogenomics analysis, capable of constructing phylogenetic networks, but its scalability in HPC environments requires attention. Similarly, [17] utilizes rooted phylogenetic network algorithms to analyze coronavirus data and presents a pipeline model for hypothesis testing, yet scalability issues in HPC settings remain unexplored.

In this work, we present the HP2NET framework [18], [19], a comprehensive approach designed to streamline the construction of phylogenetic networks within HPC computing environments and, to the best of our knowledge, it is the only study focused on evaluating the execution of such software in an HPC environment. The main contributions of this work are as follows:

- **Modeling of the HP2NET:** the framework automates data manipulation across various stages of the phylogenetic network construction, offering flexibility for deployment on both local machines and computing clusters. This versatility enables users to construct multiple networks for one or more datasets within a single execution instance, maximizing resource utilization;
- **Task packaging for pipeline execution:** through a task packaging mechanism, HP2NET efficiently manages parallelism, thereby reducing idle resources and runtime by prioritizing the execution of tasks with resolved dependencies;
- **Development of a data reuse mechanism:** HP2NET avoids the re-execution of identical tasks across different workflows, thereby enhancing efficiency;
- **Exploratory analysis of performance and scalability:** analyzing the framework’s performance to help researchers choose the most suitable approach for their needs.
- **A case study based on Dengue virus genomic data:** employed to demonstrate the practical utility and effectiveness of the HP2NET framework.

The manuscript is organized as follows. Section II introduces the background concepts relevant to the study. Section III describes the HP2NET framework and details the experimental setup. Section IV presents the results and analysis of the experiments, including the case study conducted using HP2NET. Finally, Section V summarizes the main findings and key insights derived from our research.

## II. Background

### A. Phylogenetic Trees

The evolutionary relationship among a set of individuals can be represented through a structure called a phylogenetic tree, which consists of edges, internal nodes, leaves, and, in some cases, a root. The leaves represent the individuals being studied, such as genes, species, populations, or even DNA fragments. The internal nodes correspond to hypothetical ancestors, while the edges depict the evolutionary connections between individuals. The root, when present, represents the assumed common ancestor of all individuals in the tree [20]. There are many heuristics and approaches for constructing phylogenetic trees, as well as numerous software tools that implement these methods. However, for this work, we will focus on RAxML version 8.2.12 [21], IQ-TREE version 2.2 [22], MrBayes version 3.2.7a [23], ASTRAL version 3 [24], and Quartet MaxCut version 2.10 [25], which are integrated with the HP2NET framework.

RAxML and IQ-TREE both employ maximum likelihood (ML) for constructing phylogenetic trees, generating bootstrap replicates, and implementing evolutionary models. They efficiently handle large datasets through parallel and distributed computing on HPC clusters. RAxML operates in both sequential and multi-threaded (pthreads) modes, while IQ-TREE optimizes CPU usage with the `“-T AUTO”` parameter. Both RAxML and IQ-TREE utilize branch swapping algorithms to explore alternative tree topologies by iteratively swapping branches to potentially find better-scoring configurations. These tools also employ advanced scoring functions, such as likelihood functions in ML methods, to evaluate and optimize tree topologies. They start with heuristic initial trees and iteratively refine them to improve the overall likelihood score.

MrBayes is a software for constructing phylogenetic trees using Bayesian inference, offering support for complex datasets. It provides tools for summarizing analytical results, generating consensus trees, estimating model likelihoods, and calculating posterior probabilities [23]. In combination with MBSUM and BUCKy version 1.4.4 [26], MrBayes’ output can be summarized and used to construct quartet concordance factor tables, as demonstrated in the TICR pipeline [27].

Quartet MaxCut and ASTRAL serve related purposes, as they are both used to infer species trees. However, Quartet MaxCut uses a maximum-cut approach to infer these trees from quartets, while ASTRAL infers species trees by combining gene trees.

### B. Phylogenetic Networks

Phylogenetic networks extend beyond traditional trees by representing complex scenarios like hybridization, horizontal gene transfer, and recombination, offering deeper insights into evolutionary dynamics within species or genes [28], [29]. In pathogenic organisms, these networks help unravel how genetic exchange shapes adaptation, virulence, and resistance, contributing to more effective strategies for disease surveillance, prevention, and treatment [17]. By capturing the complexity of genetic exchange, phylogenetic networks enhance our understanding of pathogen evolution.

Phylogenetic network software specializes in constructing, visualizing, and analyzing genetic data, with algorithms designed to handle reticulate events. This paper incorporates the Species Networks applying Quartets (SNaQ) algorithm [30] implemented in the PhyloNetworks package version 0.16.4 and PhyloNet version 3.8.4 [31], [32] into the architecture of HP2NET.

PhyloNet, implemented in Java, provides methods such as maximum parsimony, maximum likelihood, and maximum pseudo-likelihood for network construction based on rooted phylogenetic trees. It is used alongside the SNaQ algorithm, integrated into PhyloNetworks, which constructs phylogenetic networks from gene trees or quartet concordance factor tables, using a species tree as the initial topology. SNaQ, implemented in Julia, estimates networks using the maximum pseudo-likelihood approach and iteratively identifies the best network based on log-likelihood values. It also supports parallel execution via Julia’s distributed module, allowing computations to be distributed across multiple processes [33].

### C. Parallel Workflows Using Parsl

The Parsl library enables the creation and management of parallel workflows, allowing task execution across diverse environments such as desktop computers, clusters, clouds, and supercomputers [9].

Parsl constructs a dynamic task dependency graph, orchestrating the concurrent execution of tasks in the specified environment. By leveraging Python’s syntax, it enables parallel execution of both external programs and Python functions, using decorators like *bash apps* for external software and *python apps* for Python functions [9].

Parsl’s attributes include portability and modularization. Its programming logic and workflow execution configuration can be decoupled, supporting a myriad of architectures and configurations (named executors and execution providers). Executors oversee controlled Python task execution, while execution providers facilitate script exchanges between resources. Parsl includes several executors, including the High Throughput Executor (HTEX). The HTEX, chosen in this work for its broad applicability, implements a pilot job model and is suitable for large-scale task execution [9].

### D. Dengue Viruses (DENV)

Dengue viruses (DENV) are the primary arboviral pathogens in tropical regions globally. DENV belongs to the *Flavivirus* genus within the *Flaviviridae* family. The DENV genome encodes three structural proteins (Capsid, Membrane, Envelope) and seven non-structural proteins (NS1, NS2A, NS2B, NS3, NS4A, NS4B, NS5). DENV consists of four distinct serotypes (DENV-1 to DENV-4), each further subdivided into genotypes, associated with specific geographic regions and genetic variants [34].

## III. Methods

### A. The HP2NET Framework

#### 1) HP2NET Conceptual View

HP2NET is a high-performance computing framework designed for constructing phylogenetic trees and networks from gene sequence alignments. The conceptual view is depicted in Fig. 1. Integrated with Parsl, HP2NET seamlessly combines well-established phylogenetic software, ensuring reproducibility, efficiency, and scalability across analyses. It supports infrastructure-independent deployment, enhancing versatility and accessibility. For more details, The HP2NET framework is available on GitHub [35].

**Fig. 1.**
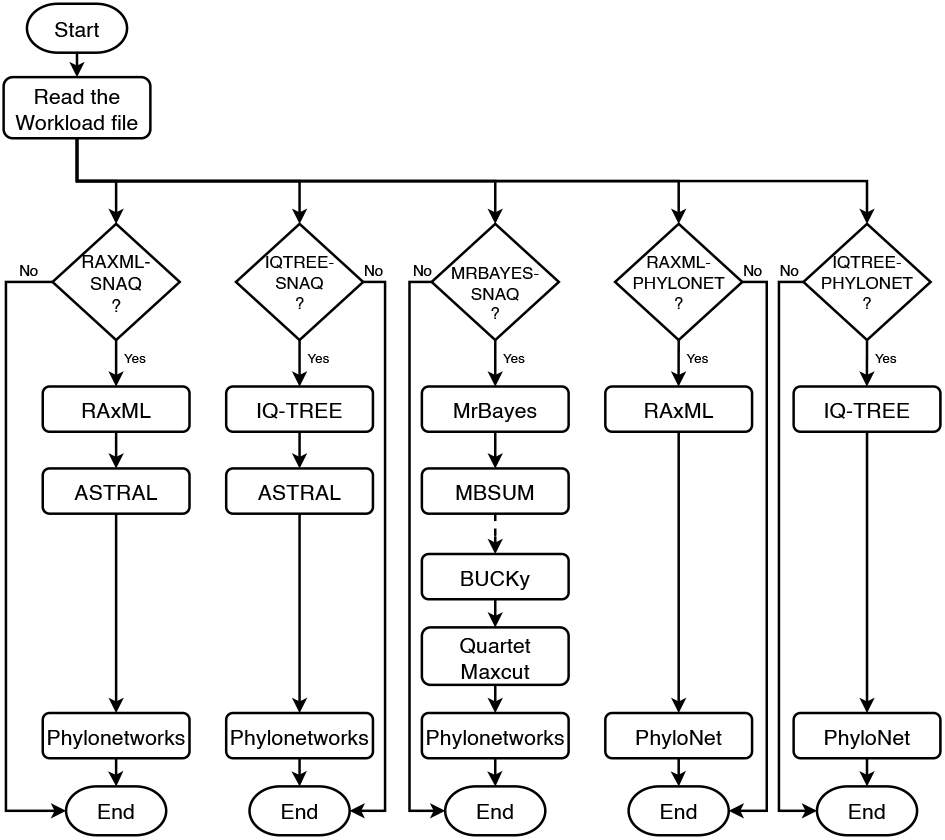
Conceptual view of HP2NET showing the five workflow configurations (RAXML-SNAQ, RAXML-PHYLONET, IQTREE-SNAQ, IQTREE-PHYLONET, MRBAYES-SNAQ). RAxML, IQ-TREE, and MrBayes run once per gene.

HP2NET offers an efficient, scalable, and flexible solution for phylogenetic network construction, achieved through five specialized workflows, referred to in this paper as RAXML-SNAQ, IQTREE-SNAQ, MRBAYES-SNAQ, RAXML-PHYLONET, and IQTREE-PHYLONET. Each workflow encompasses essential stages such as data preparation, phylogenetic tree inference, and network construction, all powered by high-performance computing and advanced workflow management systems.

### B. Parsl for Modeling HP2NET

HP2NET uses Parsl within the Python ecosystem to enhance task management and resource efficiency, similar to traditional Scientific Workflow Management Systems (SWfMS) [9]. This integration improves workflow flexibility and enables the incorporation of task packaging and data reuse mechanisms, maximizing resource utilization and reducing computation time, as detailed below. Thanks to its modular design, the framework also supports easy integration of new workflows or adaptation of existing ones. Moreover, configuring different infrastructures is straightforward.

1. **Task Packaging:** Optimizes task execution when running multiple workflows in parallel. The framework simultaneously launches all workflows and executes tasks as long as their dependencies are resolved and sufficient resources are available. In other words, tasks are executed as soon as they are ready, without waiting for other tasks to complete. Fig. 2 provides a simplified example of how tasks are scheduled using task packaging.
2. **Data Reuse:** Although Parsl includes app caching, it only stores task results after execution, which can lead to race conditions, characterizing a problem in our scenario, as race conditions could affect file creation. To address this, we implemented a caching mechanism that stores a task’s future object instead of the result, preventing duplicate executions. As a result, the framework avoids redundant tasks by executing them only once when inputs are identical across workflows. For example, the Directed Acyclic Graph (DAG) in Fig. 3 shows the concurrent execution of HP2NET’s five workflows on a 10-gene dataset. Tasks like RAxML and IQ-TREE, common to multiple workflows, run once per gene, reducing redundancy and improving makespan.

**Fig. 2.**
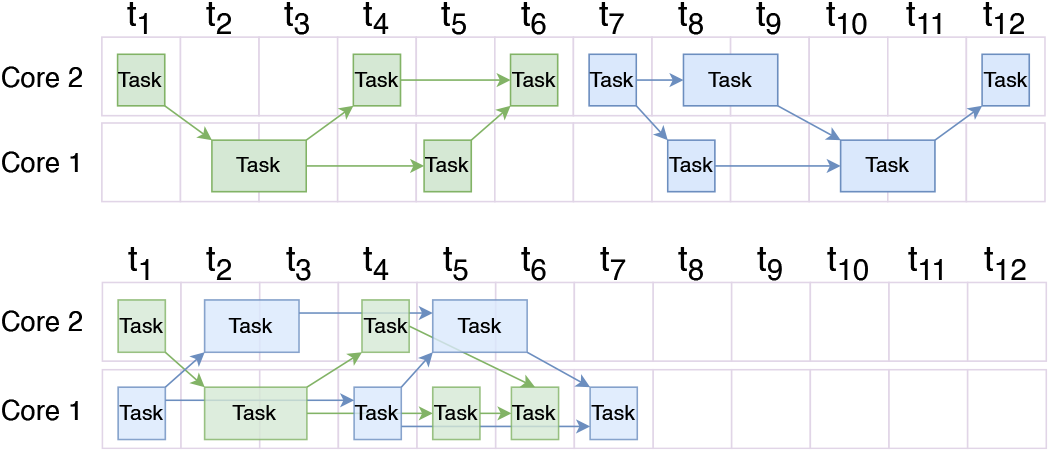
Example of task packaging: the upper image represents the execution of two pipelines (colored green and blue, respectively) sequentially without task packaging. The lower image represents the same two pipelines; however, task packaging is applied.

**Fig. 3.**
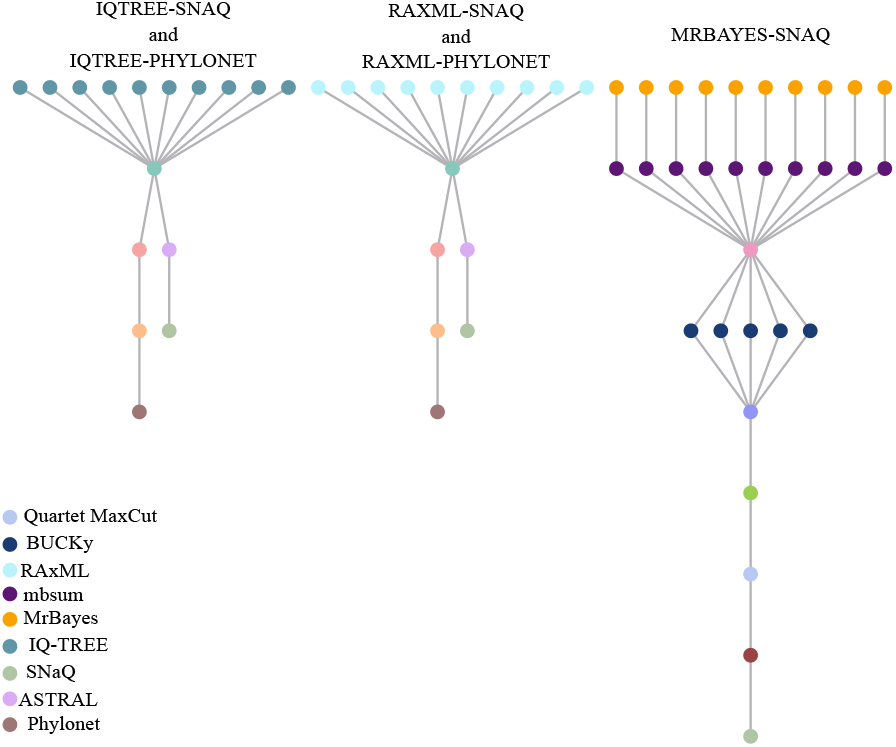
Example of data reuse in a DAG: Concurrent execution of a dataset with 10 genes across multiple workflows in HP2NET framework.

The HP2NET framework optimizes phylogenetic network construction on HPC environments by processing multiple multisequence alignments, each representing a distinct gene, alongside a JSON metadata file containing essential information like outgroup taxa. HP2NET enables users to define workloads with multiple datasets and workflows, facilitating parallel execution to maximize computational resources through data reuse and management. Ensuring reproducibility is critical in scientific experiments, and Parsl plays a pivotal role by logging and storing outputs from each app execution. This capability enables straightforward *post mortem* analysis, essential for validating and replicating experimental results.

The HP2NET framework, developed in Python v3.12.11 and powered by Parsl v1.3 for execution management, utilizes BioPython v1.75 [36] for biological data parsing and conversion. Additional functionalities are performed through scripts in other languages and specific dependencies. PhyloNetworks v0.14.3 operates with Julia v1.11.1, while ASTRAL v5.7.1 and PhyloNet v3.8.2 are executed using Java JDK v12. In our experiments, additional tasks are handled by IQ-TREE v2.2.0, RAxML v8.2.12, MrBayes v3.2.7a, Bucky v1.4.4, and Quartet Maxcut v2.10.

Installation of these tools on Linux-based systems is straightforward. Containerized versions of HP2NET are available on Docker at https://hub.docker.com/repository/docker/rafaelstjf/hp2net/general. Detailed installation instructions and troubleshooting guidance are provided on the framework’s GitHub repository.

### C. Experiments

In this section, we evaluate the performance and scalability of the HP2NET framework. HP2NET is designed to handle gene sequence alignments efficiently, scaling naturally as the number of genes increases. The framework integrates sophisticated software that supports parallelism through multi-threading and offers flexible parameter combinations, optimizing workflow efficiency.

#### 1) Environment Setup

The experiments were conducted on the Santos Dumont (SDumont) supercomputer, using a computational node equipped with two Intel Xeon Cascade Lake Gold 6252 processors (24 physical cores per socket, totaling 48 cores with hyperthreading disabled), 384 GB of RAM, and running Red Hat Enterprise Linux 8.8 with Linux Kernel version 4.18. For more detailed specifications, please visit http://sdumont.lncc.br.

#### 2) Computational Analyses of the HP2NET framework

We evaluated the performance and scalability of the HP2NET framework and identified bottlenecks within its software components using a benchmark dataset from the PhyloNetworks tutorial, available at https://github.com/crsl4/PhyloNetworks.jl/wiki/Example-Data, comprising six taxa and 100 genes, each approximately 300 base pairs long. This dataset was selected because it clearly exposes the effects of task-level parallelism while keeping the total runtime short.

1. **Performance of HP2NET and Software:** To evaluate the framework’s behavior, the five workflows were executed using a single Parsl worker, allowing only one task to run at a time. Software components that support multithreading (IQ-TREE, RAxML and SNaQ) were restricted to a single thread. After identifying bottlenecks, the multithreaded software components were analyzed individually by varying the number of threads from 1 to 24, the maximum per socket in the computational node.
2. **Scalability of HP2NET and Workflows:** Following the performance evaluation, we analyzed the scalability of HP2NET workflows by progressively increasing the number of workers from one to 48, within a single node, using the previously analyzed software configured with their optimal number of threads for the test scenario.

#### 3) Biological Analysis with HP2NET: DENV Case Study

We used HP2NET to analyze DENV-1 genomes obtained from Brazil via a GenBank [37] search conducted on June 1st, 2023, using the keywords “(complete genome dengue virus type 1) AND brazil”. After excluding sequences lacking sampling date and location information, we identified 50 genomes of DENV-1 Genotype V, ranging from 10,179 base pairs (bp) to 10,917 bp. Filtering for human hosts, we selected 43 complete genomes for genotyping, phylogenetic tree construction, and network analysis. To ensure robust statistical support, our phylogenetic analysis included an outgroup dataset comprising West Nile Virus, Zika Virus, and DENV serotypes (specifically DENV-2 through DENV-4). Our network analysis focused on Zika Virus, emphasizing its relevance and divergence.

1. **DENV-1 Genotyping Analysis:** Using the Genome Detective software, which employs Blast and phylogenetic methods, we identified the Dengue virus serotypes, genotypes, and major lineages in nucleotide sequences [38]. The genotyping analysis supported the classification of the genomes as DENV-1 Genotype V [38], [39].
2. **DENV-1 Phylogenetic Analysis:** We conducted a phylogenetic analysis of complete genome sequences of DENV-1 using RAxML version 8.2.12 [21], with the GTR+G nucleotide substitution model. To assess tree topology robustness, we used the rapid bootstrapping algorithm with 1000 replicates and tree search commands (-f a) to explore tree space. Sequences were aligned using MAFFT [40] and manually curated with Aliview [41] to remove artifacts such as insertions, deletions, and gaps. Phylogenetic tree visualization utilized Phylo.io [42] and iTOL v6 [43].
3. **DENV-1 Phylogenetic Networks:** We reduced the dataset of 43 DENV-1 genomes to 10 representative ones using the CD-HIT software [44] with a 0.99 similarity threshold to optimize computational resources in HP2NET. A Zika virus genome (accession number: MH882548) was used as an outgroup in the phylogenetic network construction process. The genomes were annoted using the FLAVi (Fast Loci Annotation of Viruses) pipeline [45] for Flaviviruses. These annotated sequences were split by genes and were aligned using MAFFT version 7.453, with the default parameters. The resulting alignments were converted into the NEXUS format, which are inputs in the HP2NET. The framework was executed using all the five workflows, enabling the construction of phylogenetic networks with Phylonet and SNaQ to detect reticulation patterns in DENV-1.

## IV. Results and Discussion

### A. Performance and Scalability of HP2NET

The runtime of each workflow executed with a single worker is shown in Fig. 4, with a breakdown of the time spent on different stages. Each stage consists of one or more executions of the same software using different (i.e., for a dataset with 100 genes, the time corresponds to running 100 independent executions of IQ-TREE). The figure highlights which stages, and consequently which software, have the greatest impact on overall execution time. The dominant stages include IQ-TREE, RAxML, SNaQ and MrBayes. From these software, RAxML, SNaQ, and IQ-TREE support internal parallelization, enabling more detailed performance analysis as the number of threads is varied, as further explored later in this section.

**Fig. 4.**
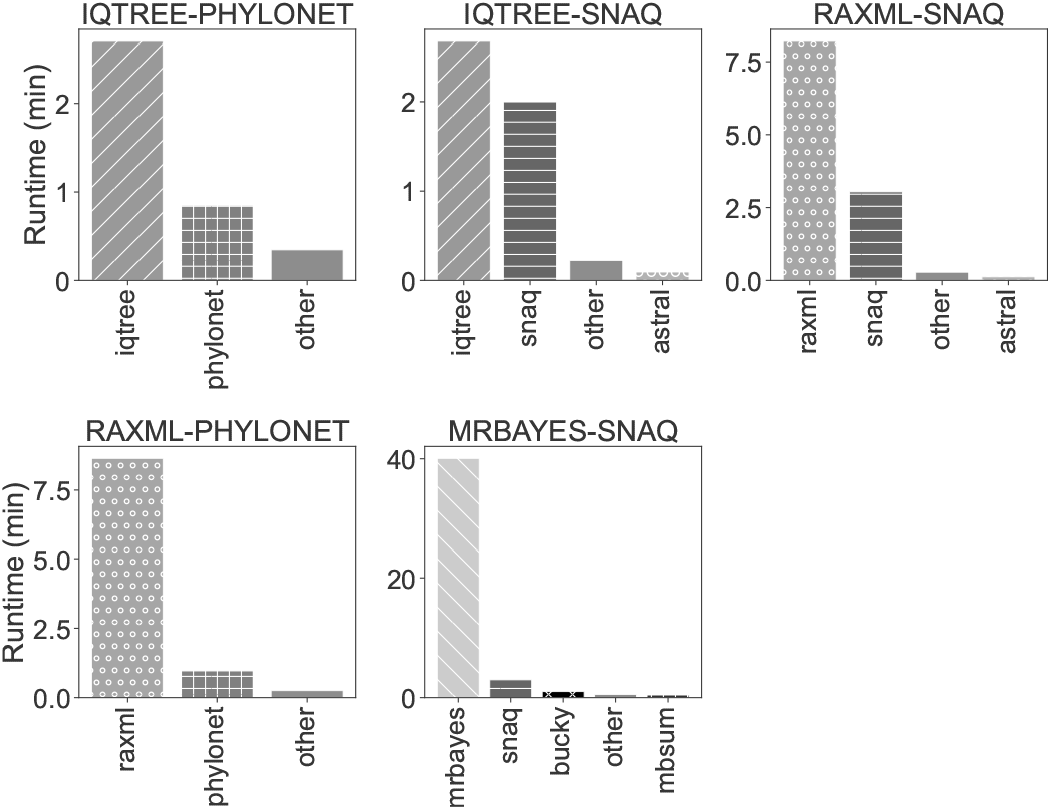
Execution time for each workflow on a dataset of 100 genes and six taxa using one Parsl worker.

### 1) Performance of theIQ-TREE Software

A comprehensive performance assessment was conducted by varying alignment lengths and the number of taxa, using IQ-TREE on three alignments: (I) 11 taxa with length 369 bp, (II) 6 taxa with 500 bp, and (III) 11 taxa with 2712 bp. The execution times for all datasets were similar to or even exceeded the sequential execution time as the number of threads increased. This phenomenon is likely due to the overhead generated by the software when parallelizing operations with short alignments. Therefore, for alignments of similar dimensions, utilizing more than one thread is unnecessary, as IQ-TREE efficiently handles the analysis sequentially and optimizes resource usage.

### 2) Performance of the RAxML Software

The three datasets previously used in IQ-TREE were also applied to RAxML.

While some datasets exhibited reduced execution times with multiple threads, the performance gain was minimal, and the additional overhead did not justify the use of parallelization. As a result, RAxML performed efficiently with the sequential version.

### 3) Performance of the SNaQ Software

Tests were conducted on two datasets to evaluate the parallel execution of the SNaQ algorithm: one with 11 taxa and 10 genes, and another with six taxa and 100 genes. Default settings were applied, including a maximum of three hybridizations and 10 runs. The results indicated that although parallelization showed some improvements, the number of threads used did not significantly affect the algorithm’s performance in this scenario.

### 4) Performance of the HP2NET Framework

Fig. 5 illustrates the average execution time for the five workflows executions and also when all of them are running concurrently in HP2NET, each repeated five times. The execution times for individual workflows exhibit similar patterns.

**Fig. 5.**
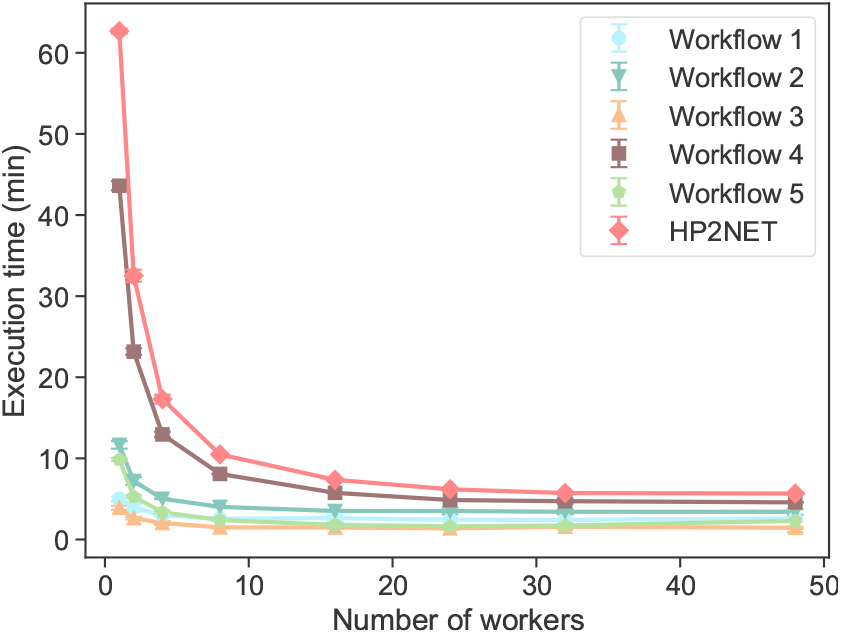
Execution time per number of workers for the five HP2NET workflows on a dataset of 100 genes and six taxa. Curves: (1) IQTREE-SNAQ, (2) RAXML-SNAQ, (3) IQTREE-PHYLONET, (4) MRBAYES-SNAQ, (5) RAXML-PHYLONET. Red line: total HP2NET execution time.

When running all five workflows concurrently, task packaging and reuse greatly improved performance, granting a 90.96% reduction in total execution time. decreasing from 62.67 to 5.67 minutes as the number of workers increased up to 48, for the tested dataset and for the used experimental environment.. This reduction was consistent across all workflows: IQTREE-SNAQ 47.86% (from 5.00 to 2.61 minutes), RAxML-SNAQ 70.69% (from 11.66 to 3.42 minutes), IQTREE-PHYLONET 62.92% (from 3.90 to 1.45 minutes), RAxML-PHYLONET 76.73% (from 9.85 to 2.29 minutes), and MRBAYES-SNAQ 89.50% (from 43.61 to 4.58 minutes). The Friedman test [46] confirmed significant differences in execution times across workers for each workflow, with all p-values lower than 3.33 × 10^−4^, indicating that performance is strongly influenced by the number of workers.

The effect of data reuse can be better observed in Fig. 6 by comparing the two lines at one worker, representing sequential execution. While the theoretical execution time for the five workflows is 74.03 minutes, HP2NET completes the execution in 62.67 minutes, achieving an approximate 15.35% reduction in execution time.

**Fig. 6.**
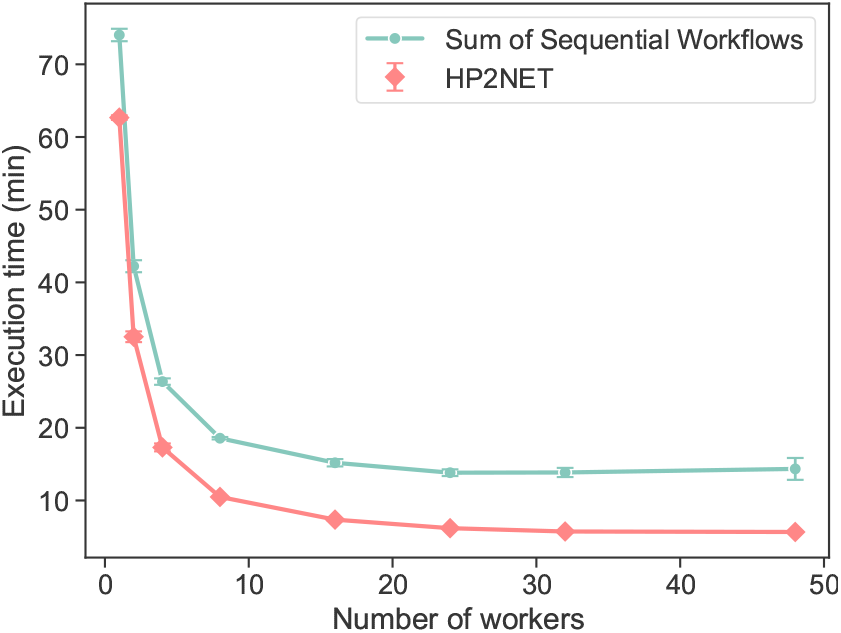
Execution time per number of workers for HP2NET on a dataset of 100 genes and six taxa. Red line: combined execution of all workflows using HP2NET; blue line: estimated sequential execution.

Fig. 7 shows the breakdown of execution stages when using 48 workers. Although the simultaneous execution of the five workflows is slower with one worker, the total time converges to that of the slowest workflow at 48 workers, with no significant difference between their execution times as indicated by a paired statistical test (*p* = 0.0625), highlighting the impact of task parallelism.

**Fig. 7.**
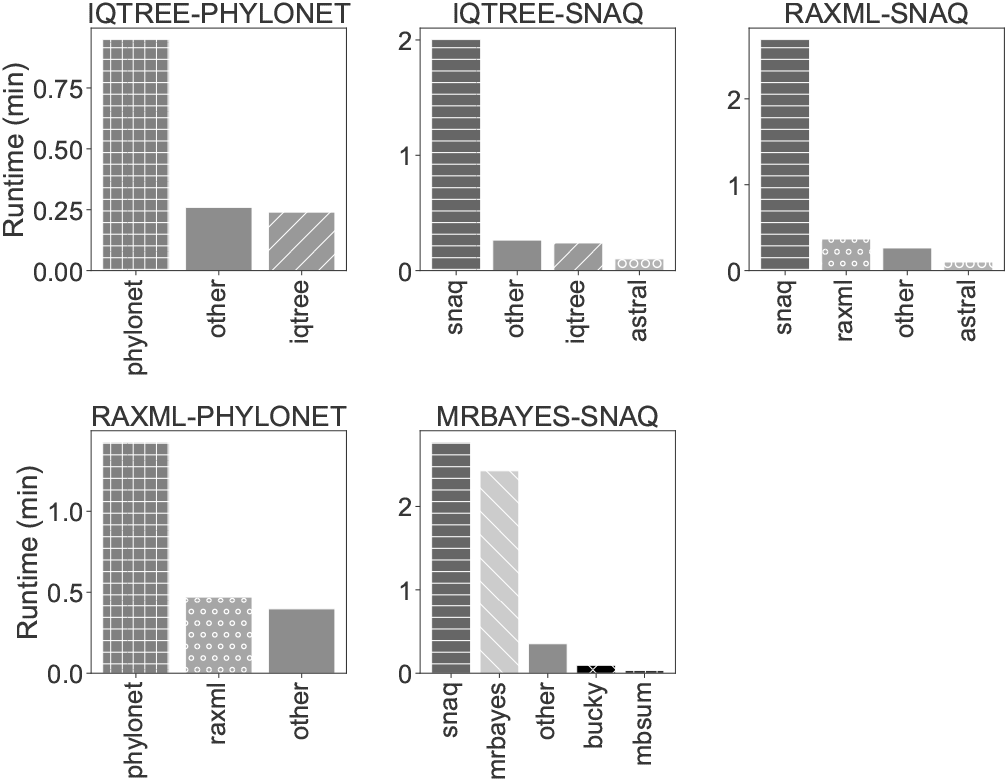
Execution time of the framework’s software when running each workflow using as input a dataset comprised of 100 genes and six taxa and 48 Parsl workers.

### B. Phylogeny and Network Analysis with HP2NET

#### 1) DENV-1 Genotyping Analysis

Our analysis of 43 complete genomic sequences of DENV-1 strains from Brazil confirms their classification as Genotype V, facilitating exploration of the reciprocal monophyly among DENV-1 genotypes. Five genotypes (I-V) of DENV-1 were previously identified using partial genomic sequences or the complete E gene [34].

Fig. 8 illustrates the phylogenetic tree with DENV-1 genotypes. The DENV-1 genomes from Brazil were identified as Genotype V (blue font/circles), revealing four main clades. Genome Detective reference sequence data are shown as follows: DENV-1 Genotype I (blue blocks/black font), DENV-1 Genotype II (red blocks/black font), DENV-1 Genotype III (yellow blocks/black font), DENV-1 Genotype IV (neon green blocks/black font), and DENV-1 Genotype V (turquoise block/black font).

**Fig. 8.**
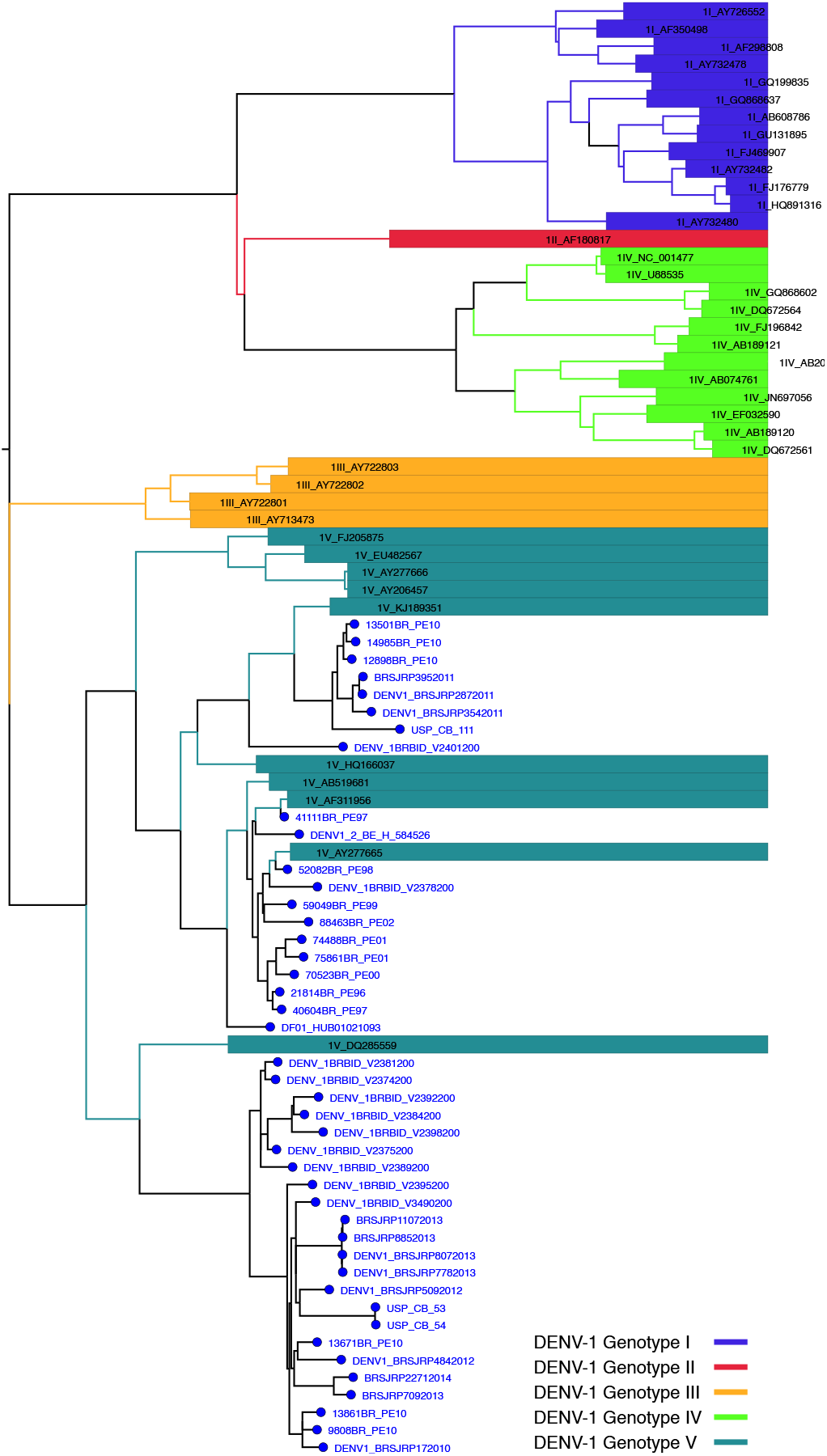
Phylogenetic Analysis of DENV-1 Genotype using Genome Detective.

#### 2) DENV-1 Phylogenetic Analysis

The sequence alignment, covering 11,208 bp and involving 43 taxa of DENV-1 Genotype V strains, alongside genomes of West Nile, Zika, and DENV serotypes (DENV-2 through DENV-4) as the outgroup, was utilized for maximum likelihood (ML) tree search. The final ML optimization likelihood was -94858.748534. Our ML tree search yielded a topology similar to that of trees generated for genotyping analysis in Genome Detective using PAUP (Fig. 8).

In this tree, the DENV-1 Genotype V tree from Brazil exhibited four main clades, with sequences from different regions from Brazil. This finding corroborates a previously reported clade shift in the DENV-1 epidemic in Brazil [47]– [50].

#### 3) DENV-1 Phylogenetic Networks

Fig. 9 displays the HP2NET networks when using all the workflows. These network topologies suggest the occurrence of hybridization events among taxa. The networks constructed using SNaQ, reticulate events involving DENV-1 ingroup taxa and the Zika outgroup were identified. Notably, sequences KP188543 and FJ850081 were implicated in reticulate events across all networks.

**Fig. 9.**
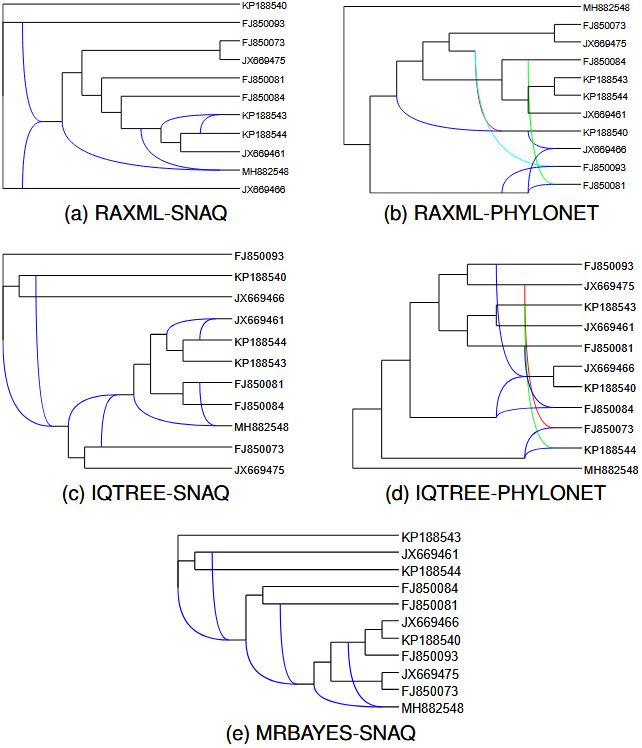
SNaQ and Phylonet networks for the dataset with 10 sequences of the DENV-1 virus genotype V. The color-coded lines (blue, red, and green) denote the same features within the networks.

## V. Conclusion

In recent years, scientific workflows have become essential tools in bioinformatics, enabling efficient orchestration of complex computational tasks within specialized High-Performance Computing (HPC) environments and Scientific Workflow Management Systems (SWfMS), enhancing resource utilization, scalability, and accelerate discoveries in bioinformatics.

This work introduces HP2NET, a versatile framework designed for constructing phylogenetic networks. Leveraging HPC capabilities, HP2NET enables parallel and distributed execution of diverse datasets and workflows. Its modular design allows for seamless addition of new components, enhancing flexibility and scalability for phylogenetic analyses.

In addition, this work conducts an initial study on Brazilian DENV-1 genomes, demonstrating genotyping and phylogenetic analysis, including phylogenetic network construction using HP2NET.

In our computational experiments, the HP2NET framework demonstrated to be a robust tool for performing phylogenetic network construction using different methodologies. The simultaneous execution of the five workflows using 48 workers, when compared to sequential execution, resulted in a reduction of up to 90.96% in total execution time, with execution times similar to the slowest execution of the isolated workflows. This reduction was a result of strategic design choices, such as data reuse, which showed a reduction of 15.35% in the execution time.

These results also highlight HP2NET’s potential for further scalability. In our experiments, its performance was constrained by the phylogenetic network construction software, which represents the final stage of the execution and provides only a single task per workflow. The experiments were executed on a single node using 48 workers, the total number of physical cores available, given the scale of the study. Nevertheless, HP2NET supports multi-node execution, which becomes more effective for larger datasets with many genes and taxa. In such scenarios, both task-level parallelism and the internal parallelism of the supporting software can be more fully exploited, an aspect that will be explored in future work. Genotyping analysis identified the genomes as genotype V, facilitating exploration of reciprocal monophyly among DENV-1 genotypes. Additionally, maximum likelihood phylogenetic tree construction indicates a clade shift in Brazil’s DENV-1 epidemic, suggesting possible reticulate events. Subsequently, five phylogenetic networks were constructed, highlighting reticulate events in the resultant networks.

In conclusion, the analysis of DENV-1 genomes reveals potential reticulate events, likely representing recombination or some form of horizontal gene transfer. However, further research is needed to confirm these findings and fully understand their implications.

## Acknowledgments

HPC resources were generously provided by the National Laboratory of Scientific Computing (LNCC/Brazil) and the Santos Dumont supercomputer. This research was funded by the National Council for Scientific and Technological Development (CNPq), specifically through the CNPq/MCTI/CT-Biotec project with Grant Number 440360/2022-6 and the University of North Carolina at Charlotte (UNC CHarlotte), Department of Bioinformatics and Genomics (BiG). Additionally, partial financial support was provided by the Coordination of Superior Level Staff Improvement (CAPES) Foundation, Brazil, under Finance Code 001.

---

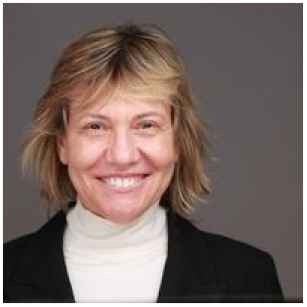

**Carla Osthoff** holds a degree in Electrical Engineering, as well as M.Sc. (1989) and Ph.D. (2000) degrees in Systems and Computer Engineering from the Federal University of Rio de Janeiro (UFRJ). Currently, she is a researcher and professor at the National Laboratory for Scientific Computing (LNCC), where she coordinates the National Center for High-Performance Processing and serves on the Santos Dumont Supercomputer Advisory Committee. Her research interests include HPC, distributed systems, parallel processing, parallel I/O systems, and scientific computing.

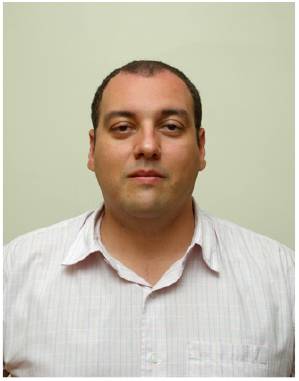

**Diego Carvalho** (M’98-SM’19) was born in Rio de Janeiro, Brazil in 1970. He received his B.S. degree in Production Engineering from UFRJ and the M.S. and Doctor’s degrees in Systems Engineering and Computer Science from PESC/COPPE. Since 2006, he has been a professor at the Department of Production Engineering of CEFET/RJ and his research interests include areas such as distributed systems, network engineering, parallel architectures, grid technologies, data mining and big data.

Dr. Carvalho is a member of the Brazilian Association of Production Engineering, Brazilian Society for the Advancement of Science, and a senior member of IEEE.

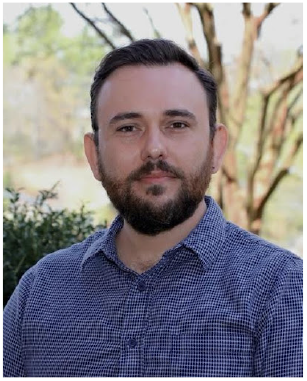

**Denis Jacob Machado** Dr. Denis Jacob Machado was born in Brazil. He earned a B.S. in Biological Sciences from São Paulo State University (UNESP) in 2009, an M.Sc. in Zoology from the University of São Paulo (USP) in 2012, and a Ph.D. in Bioinformatics from the University of São Paulo in 2018. He is an assistant professor at the University of North Carolina at Charlotte, Department of Bioinformatics and Genomics, and an early member of the Computational Intelligence to Predict Health and Environmental Risks (CIPHER) research center. He leads the Phyloinformatics Lab, focusing on pathogen evolution and biorepository data accessibility to support One Health initiatives.

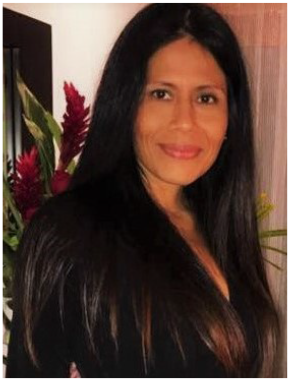

**Kary Ocaña** is a Senior Researcher at LNCC. She worked as a postdoctoral researcher at the Department of Computer Science, COPPE Institute, UFRJ, Brazil (2010-2015). Her postdoctoral fellowship was funded by FAPERJ (Postdoc Grade A) from 2013 to 2015. She received the Young Scientist of Our State award from FAPERJ (2017–2022).

She received her D.Sc. (2010) and M.Sc. (2006) in Cellular and Molecular Biology from FIOCRUZ (RJ-Brazil). Her research interests include bionformatics, scientific workflows, HPC, data analytics, and machine learning.

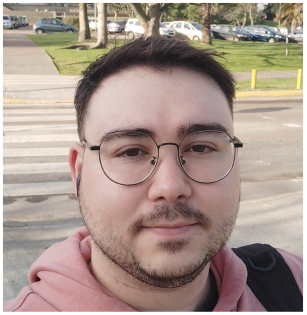

**Rafael Terra** is a Ph.D. candidate in Computational Modeling at LNCC. Master’s degree in Computing Modeling from LNCC (2022), and B.Sc in Computer Science at Federal University of Juiz de Fora (2019). His interests cover scientific workflows, HPC, and bioinformatics.

